# Antisense DNA acaricide targeting pre-rRNA of two-spotted spider mite *Tetranychus urticae* as efficacy-enhancing agent of fungus *Metarhizium robertsii*

**DOI:** 10.1101/2024.10.29.620568

**Authors:** Daria Gavrilova, Ekaterina Grizanova, Ilia Novikov, Ekaterina Laikova, Alexandra Zenkova, Vladimir Oberemok, Ivan Dubovskiy

## Abstract

Two-spotted spider mite *Tetranychus urticae* Koch (Acari: Tetranychidae) is one of the most dangerous pests in the world and one of the most pesticide-resistant species ever. Complex biological preparations are of great interest for the acaricide market because they do not poison ecosystems and do not bioaccumulate in food products, simultaneously, pests more slowly develop resistance to complex preparations. In this study we applied complex bioformulation composed of 11-mer antisense oligonucleotide (oligonucleotide acaricide or DNA acaricide) Tur-3 and fungus *Metarhizium robertsii* for *T. urticae* control. We discovered that joint contact application of DNA acaricide and fungus *M. robertsii* significantly attenuates reproduction rate of the mite. Our results indicate that DNA acaricide Tur-3 and fungus *M. robertsii* act synergistically and lead to a significant 7-times elevated mortality rate and 5-times reduced fecundity of the mite. Oligonucleotide acaricide Tur-3 causes 2,5-fold reduction of expression of the target pre-rRNA of *T. urticae* and suppresses the activity of key players of detoxifying enzymes of its defense system (phenoloxidase, esterase, glutathione-S-transferase), on average, by 2-3 times. Oligonucleotide acaricide Tur-3 interferes with protein biosynthesis causing decrease in production of defense system enzymes of the pest. Obviously, attenuation of defense system enhances fungal infection or/and fungus produces a variety of enzymes that degrade and destroy the integument structure of the pest, aiding the penetration of oligonucleotide acaricide Tur-3. This research article is the first evidence of successful application of oligonucleotide acaricide together with fungus *M. robertsii* for efficient *T. urticae* control. Combined application of oligonucleotide acaricides based on conservative antisense sequences of rDNA of pests and fungi is a potent and selective approach for eco-friendly plant protection.

## 1 Introduction

The two-spotted spider mite *Tetranychus urticae* Koch. (Acari: Tetranychidae) is a serious cosmopolitan pest of agricultural crops considered as one of the most pesticide-resistant and economically harmful species (Van Leeuwen et al., 2015). Microevolution of *T. urticae* leads to pesticide resistance and is enhanced by intensive feeding, short life cycle, high fecundity, arrhenotokous reproduction, inbreeding, and genetic mechanisms such as mutation accumulation and extensive gene copies, what pushes researchers constantly develop new acaricides. As the most polyphagous herbivore, it causes substantial crop losses, including fruits, vegetables, ornamentals. During current climate change, *T. urticae* shortens life cycle under dry and hot conditions, broadens host range, and produces more generations per year (Ximénez-Embún et al., 2017).

Mainly, chemical acaricides are used to protect crops from spider mites but the effectiveness has significantly decreased because of pesticide resistance developed by spider mites (Van Leeuwen et al., 2010). At present, spider mites demonstrate unprecedented to almost all groups of chemical plant protection products (Mosta-Sanchez, D. and John C, 2022). Thus, innovative methods for the *T. urticae* control are required for effective application in IPM programs.

One of the possible solutions to these challenges is the use of entomopathogenic fungi for biological pest control. Entomopathogenic fungi (EPF), such as *Metarhizium anisopliae, Beauveria bassiana, Isaria cateniannulata, Lecanicillium lecanii*, are commonly used to control *T. urticae* and other spider mites (Chandler et al., 2005; Maniania et al., 2008; Zenkova et al., 2020; Zhang et al., 2018). Interestingly, it has been shown that fungal infection not only leads to the death of female *T. urticae*, but also significantly reduces the fertility of surviving individuals (Shi and Feng, 2009).

The successful adaptation of these fungi in plant defense strategies necessitates a comprehensive knowledge of their ecological, evolutionary, and biological aspects, particularly their relationships with their insect hosts (Butt et al., 2016). Many defensive biochemical reactions are required by insect hosts to protect from fungal infection. The phenoloxidase system plays a crucial role in the defensive mechanisms of insects. Phenoloxidase (PO) is a copper-containing enzyme of the oxidoreductase class that oxidizes phenolic compounds. In insects, PO is localized in the hemolymph and cuticle. In insect organisms, PO exists in the form of inactive proenzymes (proPO), which are activated through limited proteolysis (Dubovskiy et al., 2016) and play an important role in cuticular protection from EPF (Dubovskiy et al., 2013a, 2013b). The primary enzymes involved in the insect detoxification process are esterases, glutathione-S-transferases (GST), and monooxygenases. These enzymes work together to help inactivate certain xenobiotics and their metabolites (Li et al., 2007). Non-specific esterases (EC 2.5.1.18) belong to the class of hydrolases that accelerate the breakdown of complex esters into alcohol and carboxylic acid. The crucial role of insect esterases are in the detoxification processes of chemical insecticides and the development of resistance to them have been demonstrated by many authors (Mamidala et al., 2011). Glutathione-S-transferase (GST) represent a group of enzymes (EC 2.5.1.18) that use reduced glutathione for conjugation with hydrophobic compounds and the reduction of organic peroxides (Snyder and Maddison, 1997). An increase in the activity and quantity of esterase and GST has been shown during the development of fungal diseases in insects (Dubovskiy et al., 2012; I.M. Dubovskiy et al., 2011; I M Dubovskiy et al., 2011; Serebrov et al., 2006).

EPF are eco-friendly biopesticides but in addition to their advantages, they also have a number of disadvantages, for example, slow action (speed of kill), which makes them generally non-preferred means of control during massive pest outbreaks, as well as the dependence of effectiveness on various biotic and abiotic factors, etc. To solve above-mentioned problems, it is proposed to use various efficacy-enhancing agents, including botanicals, bacteria (such as *Bacillus thuringiensis*, Bt), entomopathogenic nematodes, and low-dose chemical and biorational insecticides(Butt et al., 2016).

As an innovative branch of plant protection, use of contact oligonucleotide insecticides (briefly, olinscides, or DNA insecticides) against *Sternorrhyncha* members shows simplicity, flexibility, and effectiveness for control of aphids, psyllids, soft scales, armored scales, mealybugs, whiteflies, etc. (Oberemok et al., 2023a). As a novel class of insecticides, olinscides exhibit a high degree of specificity in their action, are harmless to non-target organisms, have a low environmental impact, degrade rapidly, and offer the potential for the creation of insecticides with a long-lasting effect, based on the conservative sequences of pest ribosomal RNA genes (Gal’chinsky et al., 2024, 2023; Oberemok et al., 2022, 2019; Puzanova et al., 2023). Oligonucleotide insecticides are antisense oligonucleotides (ASOs), typically 11 nt in length, that bind to complementary sites of pre-rRNAs and mature rRNAs of insect pests and degrade rRNA in rRNA–DNA hybrids recruiting RNase H (Gal’chinsky et al., 2024; Sohail et al., 2001). Oligonucleotide insecticides act through mechanism of DNA “containment”, which targets rRNA (‘arrest’ and hypercompensation), leading to degradation of target rRNA recruiting RNase H (Gal’chinsky et al., 2024). Recently, antisense oligonucleotides targeting pre-rRNA and rRNA of spider mites were proposed as oligonucleotide acaricides for *T. urticae* control (Oberemok et al., 2023a).

In medicine, ASOs represent a new and highly promising class of drugs for personalized therapies, several types of cancer, spinal muscular atrophy and dystrophy (Rinaldi and Wood, 2018; Sohail et al., 2001). It is possible to anticipate that ASO-based pesticides could become a reality in the near future, advancing by existing biotechnical solutions in plant protection. Although there is no historical precedent for an ASO approach in the realm of pesticides, ASOs for pest control are also highly appealing and effective, as they operate at the molecular level in accordance with the principle of high (Puzanova et al., 2023).

For a long time, unmodified oligodeoxyribonucleotides were believed to be rapidly cleaved by DNases and their degradation products were considered as toxic for cells (Dias and Stein, 2002). In addition, it was assumed that rRNA is resistant to degradation in the presence of antisense DNA oligonucleotides (Bachellerie et al., 2002; Will and Lührmann, 2001). The main paradigm shift with unmodified antisense DNA for plant protection was in showing that it can be a contact insecticide (Oberemok et al., 2023a). The main breakthroughs in the development of this approach were in finding the most convenient target genes (rRNA genes), showing mechanism of action (DNA containment) and in searching up insect pests (sternorrhynchans) with high susceptibility to this approach. It is important to note that if sequences from conservative parts of pest genes are used for creation of oligonucleotide pesticides, then target-site resistance to such pesticides will develop more slowly (Sharma et al., 2014). If insecticide resistance occurs, different strategies can be applied. Generally, new olinscides can be easily created displacing target site to the left or to the right from the olinscide-resistant site of target mature rRNA and/or pre-rRNA (Gal’chinsky et al., 2024). Oligonucleotide pesticides have the potential to complement the existing insecticide market and set an eco-precedent for crop protection products where the effectiveness of the insecticide will be determined by its safety for non-target organisms.

Historically, some principles of the practical application of antisense oligonucleotides for chemical reactions on biopolymers were first formulated in Novosibirsk (Russia) by N. Grineva et al. in 1967 (Belikova et al., 1967). Later, in 1978, Zamecnik and Stephenson, used modified antisense DNA against of Rous sarcoma virus in chicken embryo fibroblasts in a sequence-specific manner (Stephenson and Zamecnik, 1978). After that, the development of antisense technologies has long been primarily focused on medicine using modified antisense oligonucleotides. The mode of action of unmodified antisense DNA and its potential application as contact insecticide have not been investigated on insect pests, and no attempts were made to test this hypothesis until the beginning of the 21st century. Interestingly, though RNA interference and CRISPR/Cas9, as the next-generation technologies for plant protection, also use duplexes of unmodified nucleic acids (RNAi: guide RNA-mRNA; CRISPR/Cas9: guide RNA-genomic DNA) and nucleic acid-guided nucleases (RNAi: Argonaute, briefly Ago; CRISPR associated protein 9, briefly Cas9), nobody used another very perspective trio of participants used in ‘genetic zipper’ method: guide DNA, rRNA, and RNase (Dubovskiy et al., 2023). In the last few years, CUAD biotechnology, based on oligonucleotide pesticides, has been established as a potent and selective “genetic zipper” method against soft scale insects, armored scale insects, psyllids, mealybugs, aphids, and mites, opening new frontiers in DNA-programmable plant protection based on contact application of deoxyribonucleic acid. Already today, according to our estimations, the ‘genetic zipper’ method is potentially capable of effectively controlling 10-15% of all insect pests using a simple and flexible algorithm of DNA-programmable insect pest control.

This research article is the first evidence of joint application of an oligonucleotide acaricide and fungi *Metarhizium* spp. for efficient *T. urticae* control. Here we show positive impact from joined application of oligonucleotide acaricide (olacaricide) based on antisense rDNA and highly pathogenic strain of *Metarhizium robertsii* which synergistically complement each other’s action and provide a fundamental basis for a new approach in plant protection for mite control. It is obvious that a plethora of combinations, or repertoire, of oligonucleotide acaricides and fungal preparations will make it possible to respond very quickly to the emergence of resistance of the pest.

## 2 Materials and Methods

### 2.1 Rearing of Tetranychus urticae mite

Laboratory population of spider mite *T. urticae* was reared under laboratory conditions at Novosibirsk State Agrarian University (Novosibirsk, Russia). Mites was maintained on haricot (*Phaseolus vulgaris* L.) at 25±1°C, 50% humidity and a light:dark photoperiod (16:8 h).

### 2.2. Entomopathogenic fungus

*Metarhizium robertsii* (strain 2017) was maintained on Sabouraud dextrose agar (SDA) at 25°C for 14 days. Conidia were harvested by scraping from sporulating cultures, air-dried at room temperature (RT) for 7 days and stored at 4°C (E. V. Grizanova et al., 2019). Fungal conidia were suspended in sterile 0.03% Tween–80 and vortexed for 1 min. Viability of conidia was confirmed by incubation of these propagules on SDA and calculating germination percentage. Suspensions with at least 99% germination were used for topical infections.

### 2.3 Applied sequence of oligonucleotide acaricide Tur-3 and control oligonucleotide

We used GenBank (https://www.ncbi.nlm.nih.gov; GenBank: AM408035.1: 779-789) to design antisense oligonucleotide (oligonucleotide acaricide) Tur-3 for the experiments: 5’-AAAACATCAAG-3’ (ITS2 region of pre-rRNA). As a control we used random oligonucleotide (Alive-3): 5’-ACGTACGTACG-3’ that did not lead to the mortality of two-spotted spider mite *T. urticae* in pilot experiments (50, 100, 200 ng/μl three replicates per dose). Oligonucleotides were synthesized using DNA synthesizer, purified by reverse-phase high-performance liquid chromatography and concentrated according to previously published protocols (Oberemok et al., 2023b).

### 2.4 MALI-TOF method

A BactoSCREEN analyzer, based on a MALDI-TOF mass spectrometer, was used to assess the quality of the produced oligonucleotide acaricide and a control oligonucleotide (Litech, Moscow, Russia). The mass-to-charge ratio (m/z) of the oligonucleotides was determined as positive ions using 3-hydroxypicolinic acid as the matrix on the LaserToFLT2 Plus instrument (UK) at a 2:1 ratio. The theoretical m/z value was calculated using the ChemDraw 18.0 software, and the difference between the theoretical and experimental m/z values was no more than 10 units (ChemDraw, CambridgeSoft, USA).

### 2.5 Bioassays

The effect of the *Metarhizium robertsii* fungus and antisense oligonucleotides (ASOs) on populations of the spider mite was assessed using the plate method on plant leaves infested with the pest (Zenkova et al., 2020). The variegated red bean *Vigna angularis* plants grown on soil in a laboratory with an artificial microclimate (temperature +25±1ºC, moisture 40-60% and day light 16 hours was used. After colonization by the spider mite, the leaves were cut off and the number of spider mites on the leaf was counted with microscope. Then the leaves with the pest on them were treated using hand sprayer, with a 5 ml of suspension of the fungus water, *Metarhizium robertsii* (1×10^7^ per ml), and immediately after that with control oligonucleotide (200 ng/μl) or target oligonucleotide acaricide (200 ng/μl). Leaves were inserted in Eppendorf tube with water and placed on Petri dish at a temperature of 25°C. Counting of living spider mites were taken on the 6^th^ day after treatment.

### 2.6 Enzymes activity measurement

To estimate enzymes activity the 100 mites have been collected in 500 mkl of phosphate buffer and homogenized 1 min with ultrasonic homogenization (Q125 Sonicator, USA). The homogenates were centrifuged for 15 min, 10000 g at +4 °C. The supernatant was used for enzyme activity analysis.

Phenoloxidase (PO) activity was assayed by using a method modified from Ashida and Soderhall (Ashida and Soderhall, 1984). An aliquot of samples (10 μl) was added to microplate walls containing 200 μl of 10 mM 3,4dihydroxyphenylalanine dissolved in phosphate buffer (PBS). After 30 min, PO activity was determined by measuring the absorbance at 490 nm and 28°C with the plate reader Varioskan LUX (Thermo Fisher, USA) (Glupov et al., 2001; Grizanova et al., 2018).

Nonspecific esterase activity was estimated using p-nitrophenyl acetate hydrolysis rate following Prabhakaran et al. (Prabhakaran and Kamble, 1993). Samples (5 μl) were incubated for 10 min with 200 μl p-nitrophenyl acetate at 28°C, then, the absorbance was measured at 410 nm.

The activity of GST against 2-nitro-5-thiobenzoic acid (DNTB) was estimated by the method of Habig (Habig et al., 1974). Incubation of a 10 μl sample was performed with 1 mM glutathione and 1 mM DNTB at 25°C for 10 min. The concentration of 5-(2,4-dinitrophenyl)-glutathione was recorded at 340 nm.

PO, esterase and GST activities were converted to units of transmission density (ΔA) of the incubation mixture per min and 1 mg of protein. The protein concentration was estimated as described by Bradford (1976) with bovine serum albumin standards.

### 2.7 qRT-PCR assay

Knockdown analysis of target and control genes was quantified. Whole body samples from 50 mites were collected in 3 replicates per treatment at 24 h after inoculation with control ASOs (200 ng/mkl) and target ASOs (200 ng/mkl). Primers were designed from published *T. urticae* sequences (NCBI) or from coding sequence where high homology protein sequences could be identified from an EST library. Fore ITS2 target gene (Tur 3) (Forward: 5’-TCTGTCTGAGAGTTGAGATGTAA-3’; Reverse: 5’-CTCCTTACTAAATTGCAACGAATTG-3’). For reference genes CycA (Forward: CTTCAAGGCGGTGACTTTACC; Reverse: CCATTGAAAGGATACCTGGTCC) and Actin (Forward: GCCATCCTTCGTTTGGATTTGGCT; Reverse: TCTCGGACAATTTCTCGCTCAGCA). Primers were optimized and gene expression was measured by real-time quantitative RT-PCR according to our protocol that has been published previously (Dubovskiy et al., 2021; Grizanova et al., 2021).

### 2.7 Data analysis

Data were checked for normality (Gaussian) using the Shapiro-Wilk test. One-way parametric ANOVA with Dunnett’s multiple comparisons was applied to estimate differences in the mortality of imago and larvae and eggs of mites (n=9 per variant). One-way nonparametric ANOVA with Dunn’s multiple comparisons test was applied to estimate differences in the mortality of eggs of mites (n=9 per variant). One-way parametric ANOVA withTukey multiple comparisons test was applied to compare differences in enzymes activity (n=10-12 per variant). Data were analysed using GraphPad Prism v8.0 (GraphPad Software Inc., USA).

## 3 Results

Exposure to antisense oligonucleotide Tur-3 (ASO, or oligonucleotide acaricide) resulted in decreased expression of the target gene as well as attenuated functioning of immune and detoxifying defense systems of the mite. Infection with the fungus *Metarhizium robertsii* and treatment with ASO targeting ITS2 caused significant mortality in larvae and adults of *T. urticae*. Moreover, the combined exposure to the fungus and ASO Tur-3 led to a significant decrease in the fertility of mites.

Contact application of target ASO (Tur-3) resulted in a significant decrease in reproduction rate of the mite by two (p = 0.0156; t=3,200; df = 6.824), four (p = 0.0075; t= 4,347; df = 4.973) and six (p = 0.0157; t= 3,909; df = 4.225) times on days 5, 7 and 9, compared to the treatment with control ASO (Alive-3), respectively (Figure 1a). There was also a significant suppression of the expression level of the target gene two and a half (Actin reference gene) and two (CycA reference gene) folds compared to the treatment with the control ASO (Figure 1b).

**Figure 1.**
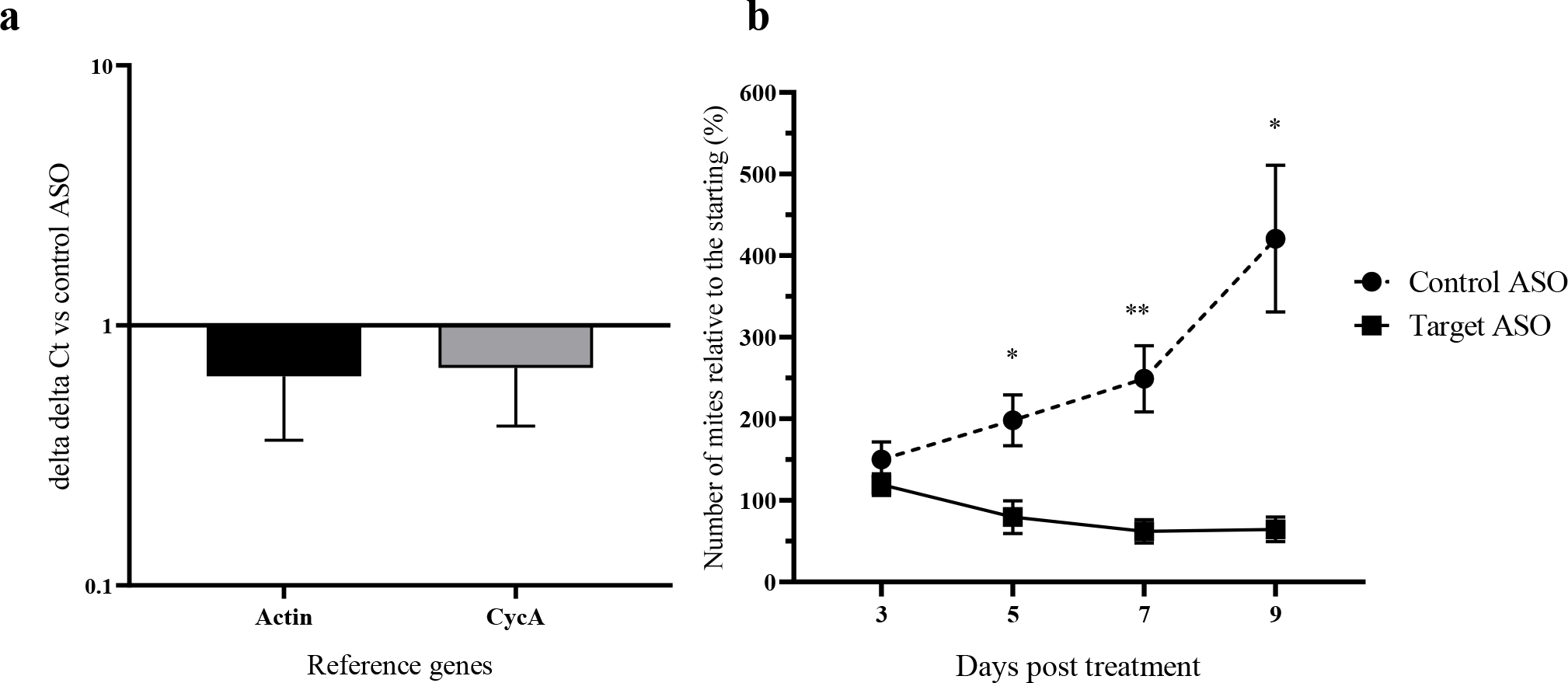
Target pre-rRNA expression on sixth day (a) and mortality (b) of spider mite *T. urticae* after treatment with control antisense oligonucleotide, Control ASO (Alive-3; 200 ng/μl) and target antisense oligonucleotide, Target ASO (Tur-3; 200 ng/μl)

Treatment of mites with target ASO Tur-3 resulted in 2-2.5-fold decrease in phenoloxidase activity compared to treatment with water (p = 0.0183 q = 4.079 df = 33) and control ASO (p = 0.0003; q= 6.320; DF=33). The activity of detoxification enzymes esterases have been decreased in mites under target ASO treatment compared with water and control ASO in 2.2 (p=0.0063; q=4.778; DF=26) and 2.5 folds (p=0.0330; q=3.783; df =26) accordingly. Activity glutathione-S-transferases also have been decreased in mites under target ASO treatments compared with water (p=0.0036; q=4.972; DF=33) and control ASO (p=0.0038; q=4.941; df =33) in 2 folds (Figure 2). The treatment of control ASO did not effect on enzyme activity compared with water treatment.

**Figure 2.**
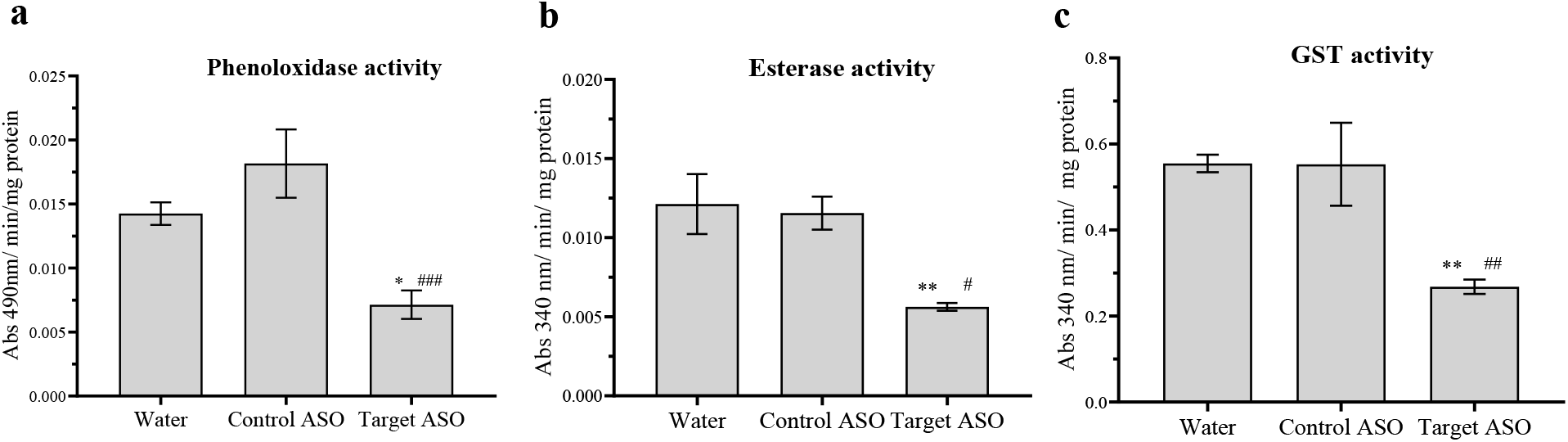
Phenoloxidase activity (a), esterase activity (b), and glutathione-S-transferase (GST) activity (c) in whole body of two-spotted spider mite *Tetranychus urticae* on sixth day after treatment with water, control antisense oligonucleotide, Control ASO (Alive-3; 200 ng/μl) and target antisense oligonucleotide, Target ASO (Tur-3; 200 ng/μl) (*p<0.05; **p<0.01; *** p<0.001 vs. water-treated Control; #p<0.05; ##p<0.01; ### p<0.001 vs. Target ASO; Tukey multiple comparisons test).

The infection with fungus *M. robertsii* reduced the number of mites larvae and adults 4.5-fold (p = < 0.0001; q=6,558; DF = 48) and target ASO 3.3-fold (p = < 0.0001; q=5,913; df = 48) on the sixth day after treatment (Fig.3). The most pronounced effect is observed in the variant of joint use of M. robertsii and target ASO Tur-3, the number of larvae and adults decreased by 7.2 times (p < 0.0001; q=7,269; DF = 48) relative to the control variant. Control ASO did not effect on survival of the mite. The mix of Control ASO with M. robertsii decreased the survival of mites 2.7-fold (p<0,0001; q=5,324; DF=48) (Figure 3).

**Figure 3.**
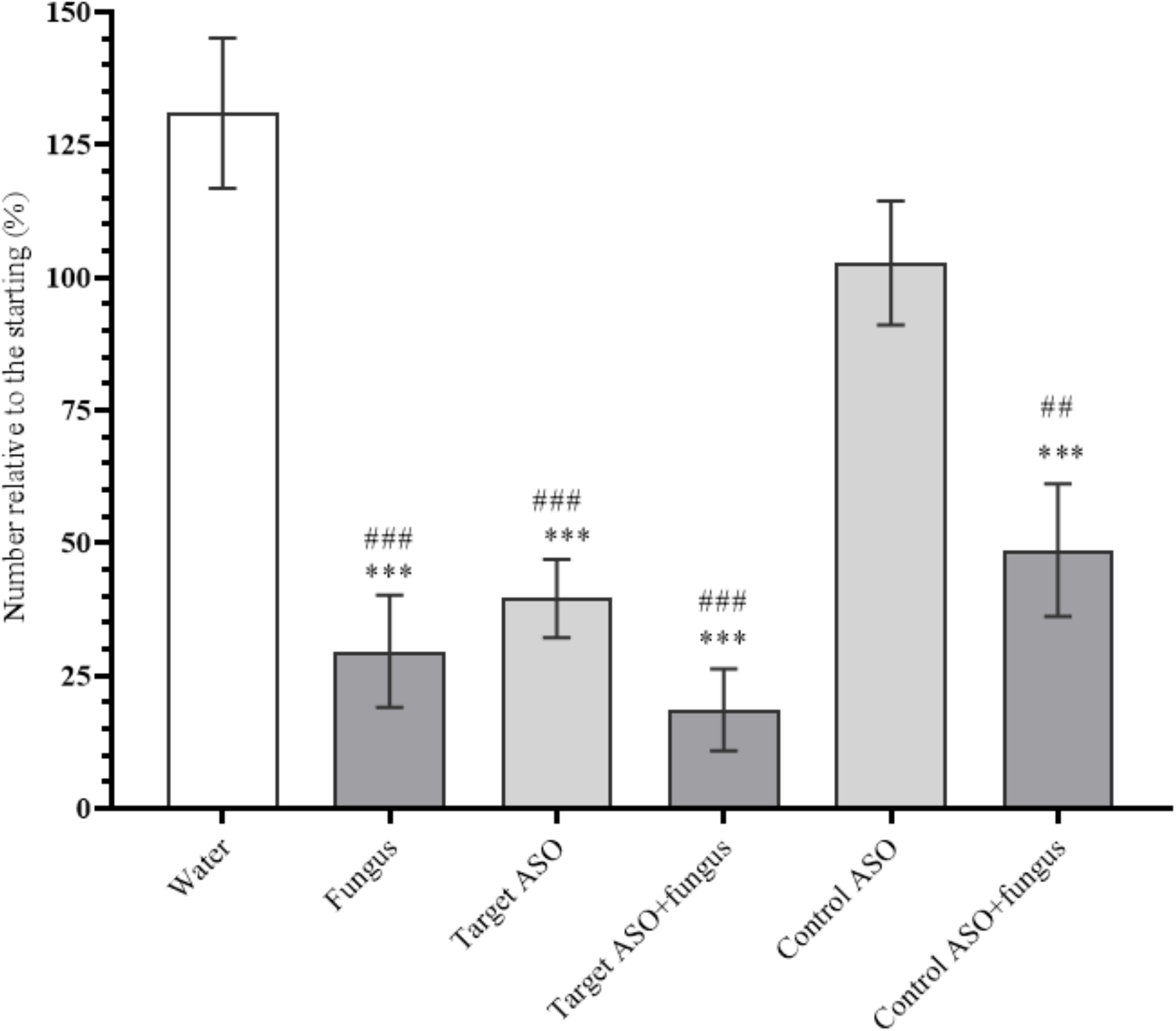
Survival rate of larvae and adults of spider mite *T. urticae* on sixth day after treatment with control antisense oligonucleotide, Control ASO (Alive-3; 200ng/l) and target antisense oligonucleotide, Target ASO (Tur-3; 200 ng/μl), fungus (*Metarhizium robertsii*; 1×10^7^) and their combinations (Control ASO+fungus, Target ASO +fungus, 10^7^+200ng/μl) (*p<0.05; **p<0.01; *** p<0.001 vs. water-treated Control, #p<0.05; ##p<0.01; ### p<0.001 vs. Control ASO; Dunnett’s multiple comparisons test).

When mites were infected with fungus or treated with target ASO, no change in fecundity was noted (Figure 4). However, with combined exposure to *M. robertsii* and target ASO, a significant decrease in mite eggs was recorded by 4.6 (p < 0.0001), 3.2 (p = 0.0183), 2.8 (p = 0.0243) fold compared with water, fungus and target ASO. Control ASO and its combination with fungus had no effect on mite fertility.

**Figure 4.**
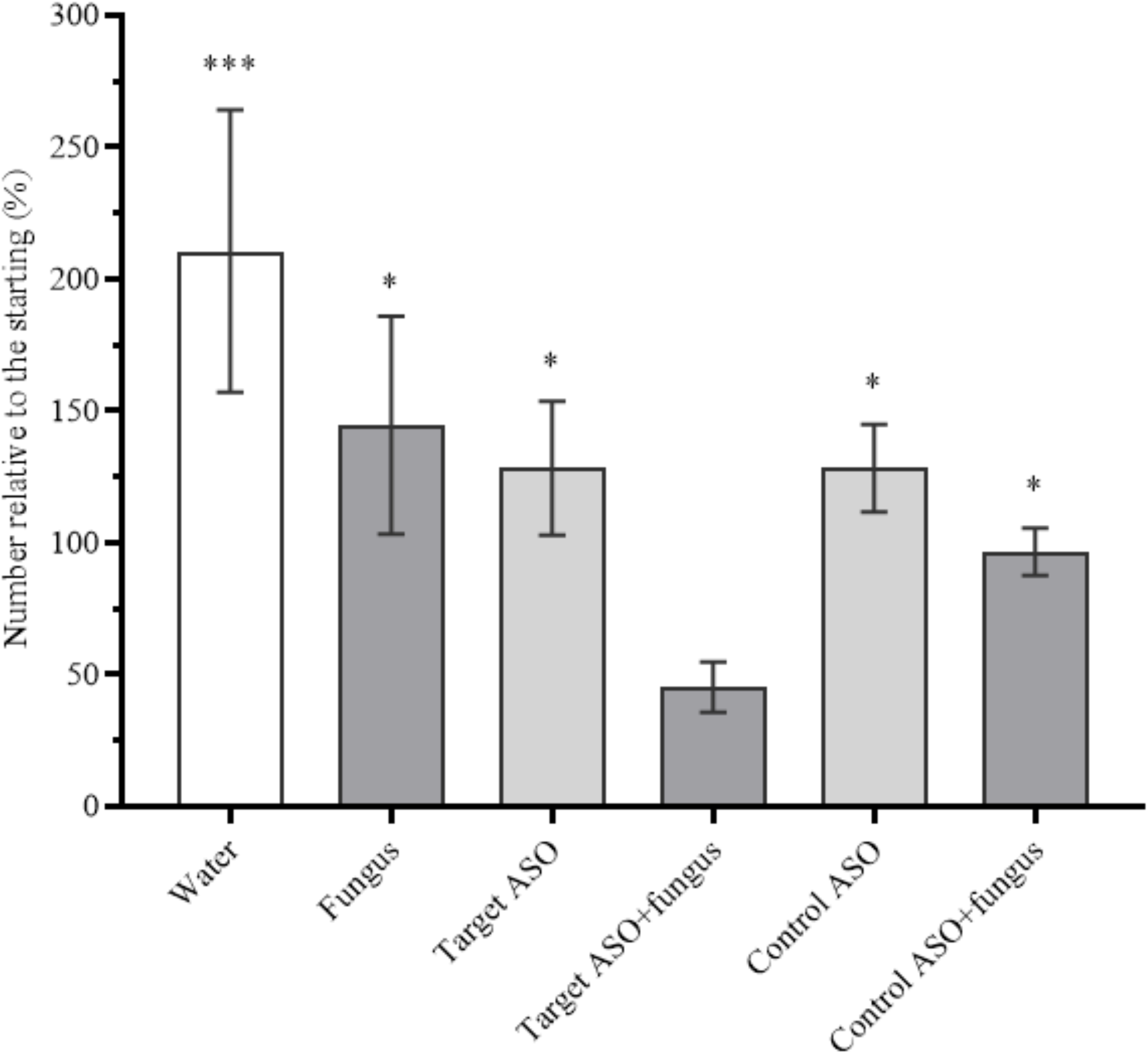
Survival rate of eggs of spider mite *T. urticae* on sixth day after treatment with control antisense oligonucleotide, Control ASO (Alive-3; 200 ng/μl) and target antisense oligonucleotide, Target ASO (Tur-3; 200 ng/μl), fungus (*Metarhizium robertsii*;10^7^) and their combinations (Control ASO+fungus, Target ASO +fungus, 10^7^+200 ng/ μl). (*p<0.05; **p<0.01; *** p<0.001 vs. Target ASO+fungus; Dunn’s multiple comparisons test).

## 4 Discussion

This study demonstrates the effectiveness of using antisense DNA (oligonucleotide acaricide) for *T. urticae* control. It was also established that the most pronounced effect is achieved when oligonucleotide acaricide is used in combination with the fungus *M. robertsii* for *T. urticae* control at all stages. The experiment revealed that oligonucleotide acaricide suppresses the activity of phenoloxidases and detoxifying enzymes, which is likely one of the main reasons for the rapid development of mycosis following the combined treatment of mites with oligonucleotide acaricide and the fungus. The combined use of fungal preparations and oligonucleotide acaricides opens up new horizons in plant protection. ITS regions of rRNA genes are moderately variable, allowing the creation of unique sequences of oligonucleotide acaricides.

In this study mite treatment with antisense DNA (ASO Tur-3) resulted in a decrease in their reproduction rate by six times because of strong acaricidal activity. Although some antisense oligos in our previous studies did not cause significant insecticidal effect on insect pests (Gal’chinsky et al., 2020; Oberemok et al., 2022, 2017). Oligonucleotide pesticides possess unprecedented selectivity in action. It should be noted that the change of one nucleotide in antisense oligos (either at 1^st^ (5′-end), 6^th^ and 11^th^ (3′-end) positions) resulted to sugnificant decrease in biological efficiency of oligonucleotide pesticides (Gal’chinsky et al., 2024; Puzanova et al., 2023). In parallel with oligonucleotide insecticides, a new class of pesticides based on dsRNA is also being developed, the action of which is based on RNAi. RNAi biocontrols has not reached substantial success in *T. urticae* control yet (Yoon et al., 2018). Oligonucleotide pesticides and RNA biocontrols, as two classes of the next-generation pesticides, are able to complement each other’s action in complex preparations for wide range of pests, including *T. urticae*, that have shown resistance to compounds from main pesticide classes (Jakubowska et al., 2022; Yoon et al., 2018).

Antisense DNA is a natural polymer that acts in a highly selective manner due to the principle of complementarity. This allows antisense DNA pesticides to combine the best features of other types of modern insecticides without the drawbacks associated with chemical insecticides (nonselective, long half-life, serious environmental consequences) or biological preparations (slow action, expensive production). Ribosomal RNA (rRNA) accounts for 80% of the cellular RNA and is more abundant and metabolically stable than other types of RNA. This makes it an excellent target for antisense DNA (Oberemok et al., 2019). Two decades ago, it was impossible to use of nucleic acids in plant protection both, conceptually and economically. When new technologies start, they often work inefficiently and are expensive; eventually they become accessible and work flawlessly. In 2008, the invention of oligonucleotide pesticides based on unmodified antisense DNA led to significant improvements in their capabilities. Today, these pesticides have remarkable characteristics: a low carbon footprint, high safety for non-target organisms, rapid biodegradation in ecosystems, and avoidance of resistance (Gal’chinsky et al., 2024).

Biopreparations for the regulation of phytophagous populations are created based on microbial pathogens and other biological agents that are elements of natural biocenoses. One of the proposed strategies for the microbiological control of spider mites involves the use of fungal entomopathogens (Zenkova et al., 2020). Entomopathogenic fungi exhibit pronounced contact action, making them effective against such sucking pests as spider mites. Several studies have already evaluated the virulence and proteolytic activities of some isolates of entomopathogenic fungi: *Metarhizium anisopliae, Beauveria bassiana, Verticillium lecanii*, and *Trichoderma harzianum*, against the spider mite. It has been found that infection of mites with the fungus *M. anisopliae* leads to significant mortality of up to 85% (Elhakim et al., 2020). Five isolates of *Metarhizium* sp. were evaluated for their pathogenicity against the spider mite and *Metarhizium* sp. resulted in the highest mortality (up to 82%) on the 5th day post-inoculation (Wasuwan et al., 2021). It was found that *T. urticae* adults were more susceptible to *M. brunneum* (some strains), *M. flavoviride* than the juvenile stages (Dogan et al., 2017). In our study, the use of the fungus *Metarhizium robertsii* also demonstrated high effectiveness, resulting in a five-fold reduction in the population at both the larval and adult stages of the mites.

To improve the effectiveness of controlling *T. urticae*, entomopathogenic fungi can be combined with other biological control agents or low doses of microbial metabolites. For instance, some researchers suggest using a combination of entomopathogenic fungi and avermectin metabolites to combat mosquitoes (Noskov et al., 2019) and the Colorado potato beetle (Tomilova et al., 2016). It has been established that among the reasons for the synergistic effect may be the disruption of immune and detoxifying systems under combined exposure (Butt et al., 2016). In our study, we also examined the impact of antisense oligonucleotides on the immune and detoxifying system of the common spider mite *Tetranychus urticae*. When *T. urticae* was exposed to unmodified antisense oligonucleotides, we observed a reduction in phenoloxidase activity, one of the key enzymes of cellular and humoral immunity (Dubovskiy et al., 2016). Phenoloxidase (PO) activity, is clearly correlated with increased fungal *Beauveria bassiana* resistance of the wax moth *Galleria mellonella* (Dubovskiy et al., 2013a). Moreover, population of the wax moth *G. mellonella* under constant selective pressure from the pathogenic fungus *B. bassiana* exhibit significantly enhanced resistance by prioritizing and re-allocating pathogen-species-specific augmentations (cuticular phenoloxidase activity as well) to integumental front-line defenses (Dubovskiy et al., 2013a; Dubovskiy et al., 2013). The changes in the immune responses of the Colorado potato beetle larvae, especially in PO activity in the hemolymph and cuticle, correlate with strain-specific virulence of different entomopathogenic fungi *Metarhizium* species (Tyurin et al., 2016). Additionally, esterase activity and glutathione S-transferase activity, essential enzymes of the detoxifying system responsible for detoxifying endogenous and exogenous toxins have been demonstrated as important parts of protection during insects infections (E.V. Grizanova et al., 2019; Yaroslavtseva et al., 2017). Altogether, the suppression of the mites’ immune and detoxifying systems by oligonucleotide acaricide can significantly reduce resistance to fungal infection, as established under the combined effect of the fungus and antisense DNA.

To prevent the reinfestation of plants by the spider mite, it is essential to target the egg stage of the pest, as it is the most challenging phase to control due to its resilience (Ximénez-Embún et al., 2017). In this study, it was also experimentally proven that the combined effect of the fungus and antisense oligonucleotide resulted in a significant reduction in mite fertility. The mechanism of this action is not yet fully understood and requires further investigation.

## 5. Conclusions

The adaptation of oligonucleotide pesticides into plant protection practice will help create a flexible system for pesticide production to the constantly changing genetics of pests during microevolution. Joint application of fungal preparations and oligonucleotide pesticides can solve pesticide resistance problem for particular pests and create formulations with multidecade utility. So far, oligonucleotide pesticides have enjoyed substantial success against sternorrhynchans (Hemiptera) and spider mites but more complex formulations of oligonucleotide pesticides with fungi and auxiliary substances (spreaders, adhesives, penetrators, etc.) may be effective in controlling other pests. The balance between the price of such complex pesticides (oligonucleotide acaricides + fungal preparations) and the effectiveness of the formulations can ensure the popularity of such preparations and may lead to high crop yields without risks for human health and the environment.

## Author Contributions

Conceptualization, ID, EG and VO; methodology, DG, EG, IN, EL; formal analysis, DG, ID; investigation, DG, EG, ID; resources, AZ, ID; data curation, DG, AZ; writing—original draft preparation, DG, EG, VO, ID; funding acquisition, VO, ID. All authors have read and agreed to the published version of the manuscript.

## Institutional Review Board Statement

Not applicable.

## Informed Consent Statement

Not applicable.

## Data Availability Statement

Data will be made available on request.

## References

Ashida, M., Soderhall, K., 1984. The prophenoloxidase activating system in Crayfish. Comparative Biochemistry and Physiology B-Biochemistry & Molecular Biology 77, 21–26. 10.1016/0305-0491(84)90217-7

Bachellerie, J.P., Cavaillé, J., Hüttenhofer, A., 2002. The expanding snoRNA world. Biochimie. 10.1016/S0300-9084(02)01402-5

Belikova, A.M., Zarytova, V.F., Grineva, N.I., 1967. Synthesis of ribonucleosides and diribonucleoside phosphates containing 2-chloro-ethylamine and nitrogen mustard residues. Tetrahedron Lett 8, 3557–3562. 10.1016/S0040-4039(01)89794-X

Butt, T.M., Coates, C.J., Dubovskiy, I.M., Ratcliffe, N.A., 2016. Entomopathogenic Fungi: New Insights into Host-Pathogen Interactions. Adv Genet 94, 307–364. 10.1016/bs.adgen.2016.01.006

Chandler, D., Davidson, G., Jacobson, R.J., 2005. Laboratory and glasshouse evaluation of entomopathogenic fungi against the two-spotted spider mite, Tetranychus urticae (Acari: Tetranychidae), on tomato, Lycopersicon esculentum. Biocontrol Sci Technol. 10.1080/09583150410001720617

Dias, N., Stein, C.A., 2002. Antisense oligonucleotides: Basic concepts and mechanisms. Mol Cancer Ther.

Dogan, Y.O., Hazir, S., Yildiz, A., Butt, T.M., Cakmak, I., 2017. Evaluation of entomopathogenic fungi for the control of Tetranychus urticae (Acari: Tetranychidae) and the effect of Metarhizium brunneum on the predatory mites (Acari: Phytoseiidae). Biological Control 111, 6–12. 10.1016/j.biocontrol.2017.05.001

Dubovskiy, I.M., Grizanova, E.V., Gerasimova, S.V., 2023. Plant recombinant gene technology for pest control in XXI century: from simple transgenesis to CRISPR/Cas, in: Kumar, A., Arora, S., Ogita, S., Yau, Y.-Y., Mukherjee, K. (Eds.), Gene Editing in Plants: CRISPR-Cas and Its Applications. Springer Nature Singapore Pte Ltd., Singapore.

Dubovskiy, I M, Grizanova, E. V, Ershova, N.S., Rantala, M.J., Glupov, V. V, 2011. The effects of dietary nickel on the detoxification enzymes, innate immunity and resistance to the fungus Beauveria bassiana in the larvae of the greater wax moth Galleria mellonella. Chemosphere 85, 92–96. 10.1016/j.chemosphere.2011.05.039

Dubovskiy, I.M., Grizanova, E. V., Tereshchenko, D., Krytsyna, T.I., Alikina, T., Kalmykova, G., Kabilov, M., Coates, C.J., 2021. Bacillus thuringiensis Spores and Cry3A Toxins Act Synergistically to Expedite Colorado Potato Beetle Mortality. Toxins (Basel) 13, 746. 10.3390/toxins13110746

Dubovskiy, I.M., Kryukova, N.A., Glupov, V.V., Ratcliffe, N.A., 2016. Encapsulation and nodulation in insects. Invertebrate Survival Journal 13.

Dubovskiy, I.M., Slyamova, N.D., Kryukov, V.Y., Yaroslavtseva, O.N., Levchenko, M.V., Belgibaeva, A.B., Adilkhankyzy, A., Glupov, V.V., 2012. The activity of nonspecific esterases and glutathione-S-transferase in Locusta migratoria larvae infected with the fungus Metarhizium anisopliae (Ascomycota, Hypocreales). Entomol Rev 92. 10.1134/S0013873812010022

Dubovskiy, I.M., Slyamova, N.D., Yu Kryukov, V., Yaroslavtseva, O.N., Levchenko, M.V., Belgibaeva, A.B., Adilkhankyzy, A., Glupov, V.V., 2011. Activity of nonspecific esterases and glutathione S-transferase of locusta migratoria larvae in infection with metarhizium anisopliae fungus (Ascomycota, hypocreales). Zool Zhurnal 90.

Dubovskiy, I.M., Whitten, M.A., Kryukov, V.Y., Yaroslavtseva, O.N., Grizanova, E. V., Greig, C., Mukherjee, K., Vilcinskas, A., Mitkovets, P. V., Glupov, V. V., Butt, T.M., 2013a. More than a colour change: insect melanism, disease resistance and fecundity. Proceedings of the Royal Society B-Biological Sciences 280, 10.

Dubovskiy, I.M., Whitten, M.M.A., Yaroslavtseva, O.N., Greig, C., Kryukov, V.Y., Grizanova, E. V., Mukherjee, K., Vilcinskas, A., Glupov, V. V., Butt, T.M., 2013b. Can Insects Develop Resistance to Insect Pathogenic Fungi? PLoS One 8.

Elhakim, E., Mohamed, O., Elazouni, I., 2020. Virulence and proteolytic activity of entomopathogenic fungi against the two-spotted spider mite, Tetranychus urticae Koch (Acari: Tetranychidae). Egypt J Biol Pest Control 30, 30. 10.1186/s41938-020-00227-y

Gal’chinsky, N., Useinov, R., Yatskova, E., Laikova, K., Novikov, I., Gorlov, M., Trikoz, N., Sharmagiy, A., Plugatar, Y., Oberemok, V., 2020. A breakthrough in the efficiency of contact DNA insecticides: Rapid high mortality rates in the sap-sucking insects Dynaspidiotus britannicus Comstock and Unaspis euonymi Newstead. J Plant Prot Res 60. 10.24425/jppr.2020.133315

Gal’chinsky, N. V., Yatskova, E. V., Novikov, I.A., Sharmagiy, A.K., Plugatar, Y. V., Oberemok, V. V., 2024. Mixed insect pest populations of Diaspididae species under control of oligonucleotide insecticides: 3′-end nucleotide matters. Pestic Biochem Physiol 200. 10.1016/j.pestbp.2024.105838

Gal’chinsky, N. V., Yatskova, E. V., Novikov, I.A., Useinov, R.Z., Kouakou, N.J., Kouame, K.F., Kra, K.D., Sharmagiy, A.K., Plugatar, Y. V., Laikova, K. V., Oberemok, V. V., 2023. Icerya purchasi Maskell (Hemiptera: Monophlebidae) Control Using Low Carbon Footprint Oligonucleotide Insecticides. Int J Mol Sci 24. 10.3390/ijms241411650

Glupov, V. V., Khvoshevskaya, M.F., Lozinskaya, Y.L., Dubovski, I.M., Martemyanov, V. V., Sokolova, J.Y., 2001. Application of the nitroblue tetrazolium-reduction method for studies on the production of reactive oxygen species in insect haemocytes. Cytobios 413, 165–178.

Grizanova, E.V., Krytsyna, T.I., Surcova, V.S., Dubovskiy, I.M., 2019. The role of midgut nonspecific esterase in the susceptibility of Galleria mellonella larvae to Bacillus thuringiensis. J Invertebr Pathol 166. 10.1016/j.jip.2019.107208

Grizanova, E.V., Semenova, A.D., Komarov, D.A., Chertkova, E.A., Slepneva, I.A., Dubovskiy, I.M., 2018. Maintenance of redox balance by antioxidants in hemolymph of the greater wax moth Galleria mellonella larvae during encapsulation response. Arch Insect Biochem Physiol. 10.1002/arch.21460

Grizanova, E. V., Coates, C.J., Dubovskiy, I.M., Butt, T.M., 2019. Metarhizium brunneum infection dynamics differ at the cuticle interface of susceptible and tolerant morphs of Galleria mellonella. Virulence. 10.1080/21505594.2019.1693230

Grizanova, Ekaterina. V., Coates, Christopher.J., Butt, Tariq.M., Dubovskiy, Ivan.M., 2021. RNAi-mediated suppression of insect metalloprotease inhibitor (IMPI) enhances Galleria mellonella susceptibility to fungal infection. Dev Comp Immunol 122, 104126. 10.1016/J.DCI.2021.104126

Habig, W.H., Pabst, M.J., Jakoby, W.B., 1974. Glutathione S transferases. The first enzymatic step in mercapturic acid formation. Journal of Biological Chemistry 249. 10.1016/S0021-9258(19)42083-8

Jakubowska, M., Dobosz, R., Zawada, D., Kowalska, J., 2022. A Review of Crop Protection Methods against the Twospotted Spider Mite—Tetranychus urticae Koch (Acari: Tetranychidae)—with Special Reference to Alternative Methods. Agriculture (Switzerland). 10.3390/agriculture12070898

Li, X., Schuler, M.A., Berenbaum, M.R., 2007. Molecular Mechanisms of Metabolic Resistance to Synthetic and Natural Xenobiotics. Annu Rev Entomol. 10.1146/annurev.ento.51.110104.151104

Mamidala, P., Jones, S.C., Mittapalli, O., 2011. Metabolic resistance in bed bugs. Insects. 10.3390/in-sects2010036

Maniania, N.K., Bugeme, D.M., Wekesa, V.W., Delalibera, I., Knapp, M., 2008. Role of entomopathogenic fungi in the control of Tetranychus evansi and Tetranychus urticae (Acari: Tetranychidae), pests of horticultural crops. Exp Appl Acarol. 10.1007/s10493-008-9180-8

Mosta-Sanchez, D. and John C, W., 2022. Arthropods resistant to pesticides database [WWW Document]. https://www.pesticideresistance.org.

Noskov, Y.A., Polenogova, O. V., Yaroslavtseva, O.N., Belevich, O.E., Yurchenko, Y.A., Chertkova, E.A., Kryukova, N.A., Kryukov, V.Y., Glupov, V. V., 2019. Combined effect of the entomopathogenic fungus Metarhizium robertsii and avermectins on the survival and immune response of Aedes aegypti larvae. PeerJ 2019. 10.7717/peerj.7931

Oberemok, V. V., Gal’chinsky, N. V., Useinov, R.Z., Novikov, I.A., Puzanova, Y. V., Filatov, R.I., Kouakou, N.J., Kouame, K.F., Kra, K.D., Laikova, K. V., 2023a. Four Most Pathogenic Superfamilies of Insect Pests of Suborder Sternorrhyncha: Invisible Superplunderers of Plant Vitality. Insects. 10.3390/in-sects14050462

Oberemok, V. V., Laikova, K. V., Gal’chinsky, N. V., Useinov, R.Z., Novikov, I.A., Temirova, Z.Z., Shumskykh, M.N., Krasnodubets, A.M., Repetskaya, A.I., Dyadichev, V. V., Fomochkina, I.I., Bessalova, E.Y., Makalish, T.P., Gninenko, Y.I., Kubyshkin, A. V., 2019. DNA insecticide developed from the Lymantria dispar 5.8S ribosomal RNA gene provides a novel biotechnology for plant protection. Sci Rep 9, 6197. 10.1038/s41598-019-42688-8

Oberemok, V. V., Laikova, K. V., Yurchenko, K.A., Novikov, I.A., Makalish, T.P., Kubyshkin, A. V., Andreeva, O.A., Bilyk, A.I., 2023b. Adjuvant Oligonucleotide Vaccine Increases Survival and Improves Lung Tissue Condition of B6.Cg-Tg (K18-ACE2)2 Transgenic Mice. Sci Pharm 91. 10.3390/scipharm91030035

Oberemok, V. V., Laikova, K. V., Zaitsev, A.S., Temirova, Z.Z., Gal’chinsky, N. V., Nyadar, P.M., Shumskykh, M.N., Zubarev, I. V., 2017. The need for the application of modern chemical insecticides and environmental consequences of their use: A mini review. J Plant Prot Res. 10.1515/jppr-2017-0044

Oberemok, V. V., Useinov, R.Z., Skorokhod, O.A., Gal’chinsky, N. V., Novikov, I.A., Makalish, T.P., Yatskova, E. V., Sharmagiy, A.K., Golovkin, I.O., Gninenko, Y.I., Puzanova, Y. V., Andreeva, O.A., Alieva, E.E., Eken, E., Laikova, K. V., Plugatar, Y. V., 2022. Oligonucleotide Insecticides for Green Agriculture: Regulatory Role of Contact DNA in Plant–Insect Interactions. Int J Mol Sci 23. 10.3390/ijms232415681

Prabhakaran, S.K., Kamble, S.T., 1993. Activity and electrophoretic characterization of esterases in insecticide-resistant and susceptible strains of German cockroach (Dictyoptera: Blattellidae). J Econ Entomol. 10.1093/jee/86.4.1009

Puzanova, Y. V., Novikov, I.A., Bilyk, A.I., Sharmagiy, A.K., Plugatar, Y. V., Oberemok, V. V., 2023. Perfect Complementarity Mechanism for Aphid Control: Oligonucleotide Insecticide Macsan-11 Selectively Causes High Mortality Rate for Macrosiphoniella sanborni Gillette. Int J Mol Sci 24. 10.3390/ijms241411690

Rinaldi, C., Wood, M.J.A., 2018. Antisense oligonucleotides: The next frontier for treatment of neurological disorders. Nat Rev Neurol. 10.1038/nrneurol.2017.148

Serebrov, V. V., Gerber, O.N., Malyarchuk, A.A., Martemyanov, V. V., Alekseev, A.A., Glupov, V. V., 2006. Effect of entomopathogenic fungi on detoxification enzyme activity in greater wax moth Galleria mellonella L. (Lepidoptera, Pyralidae) and role of detoxification enzymes in development of insect resistance to entomopathogenic fungi. Biology Bulletin 33, 581–586. 10.1134/S1062359006060082

Sharma, V.K., Sharma, R.K., Singh, S.K., 2014. Antisense oligonucleotides: Modifications and clinical trials. Medchemcomm. 10.1039/c4md00184b

Shi, W.B., Feng, M.G., 2009. Effect of fungal infection on reproductive potential and survival time of Tetranychus urticae (Acari: Tetranychidae). Exp Appl Acarol. 10.1007/s10493-009-9238-2

Snyder, M.J., Maddison, D.R., 1997. Molecular phylogeny of glutathione-S-transferases. DNA Cell Biol 16. 10.1089/dna.1997.16.1373

Sohail, M., Hochegger, H., Klotzbücher, A., Le Guellec, R., Hunt, T., Southern, E.M., 2001. Antisense oligonucleotides selected by hybridisation to scanning arrays are effective reagents in vivo. Nucleic Acids Res 29. 10.1093/nar/29.10.2041

Stephenson, M.L., Zamecnik, P.C., 1978. Inhibition of Rous sarcoma viral RNA translation by a specific oligodeoxyribonucleotide. Proc Natl Acad Sci U S A 75. 10.1073/pnas.75.1.285

Tomilova, O.G., Kryukov, V.Y., Duisembekov, B.A., Yaroslavtseva, O.N., Tyurin, M. V., Kryukova, N.A., Skorokhod, V., Dubovskiy, I.M., Glupov, V. V., 2016. Immune-physiological aspects of synergy between avermectins and the entomopathogenic fungus Metarhizium robertsii in Colorado potato beetle larvae. J Invertebr Pathol. 10.1016/j.jip.2016.08.008

Van Leeuwen, T., Tirry, L., Yamamoto, A., Nauen, R., Dermauw, W., 2015. The economic importance of acaricides in the control of phytophagous mites and an update on recent acaricide mode of action research. Pestic Biochem Physiol. 10.1016/j.pestbp.2014.12.009

Van Leeuwen, T., Vontas, J., Tsagkarakou, A., Dermauw, W., Tirry, L., 2010. Acaricide resistance mechanisms in the two-spotted spider mite Tetranychus urticae and other important Acari: A review. Insect Biochem Mol Biol. 10.1016/j.ibmb.2010.05.008

Wasuwan, R., Phosrithong, N., Promdonkoy, B., Sangsrakru, D., Sonthirod, C., Tangphatsornruang, S., Likhitrattanapisal, S., Ingsriswang, S., Srisuksam, C., Klamchao, K., Suksangpanomrung, M., Hleepongpanich, T., Reungpatthanaphong, S., Tanticharoen, M., Amnuaykanjanasin, A., 2021. The Fungus Metarhizium sp. BCC 4849 Is an Effective and Safe Mycoinsecticide for the Management of Spider Mites and Other Insect Pests. Insects 13, 42. 10.3390/insects13010042

Will, C.L., Lührmann, R., 2001. Spliceosomal UsnRNP biogenesis, structure and function. Curr Opin Cell Biol. 10.1016/S0955-0674(00)00211-8

Ximénez-Embún, M.G., Castañera, P., Ortego, F., 2017. Drought stress in tomato increases the performance of adapted and non-adapted strains of Tetranychus urticae. J Insect Physiol 96. 10.1016/j.jinsphys.2016.10.015

Yaroslavtseva, O.N., Dubovskiy, I.M., Khodyrev, V.P., Duisembekov, B.A., Kryukov, V.Y., Glupov, V.V., 2017. Immunological mechanisms of synergy between fungus Metarhizium robertsii and bacteria Bacillus thuringiensis ssp. morrisoni on Colorado potato beetle larvae. J Insect Physiol 96. 10.1016/j.jin-sphys.2016.10.004

Yoon, J.S., Sahoo, D.K., Maiti, I.B., Palli, S.R., 2018. Identification of target genes for RNAi-mediated control of the Twospotted Spider Mite. Sci Rep 8. 10.1038/s41598-018-32742-2

Zenkova, A.A., Grizanova, E. V., Andreeva, I. V., Gerne, D.Y., Shatalova, E.I., Cvetcova, V.P., Dubovskiy, I.M., 2020. Effect of fungus Lecanicillium lecanii and bacteria Bacillus thuringiensis, Streptomyces avermitilis on two-spotted spider mite Tetranychus urticae (Acari: Tetranychidae) and predatory mite Phytoseiulus persimilis (Acari: Phytoseiidae). J Plant Prot Res 60. 10.24425/jppr.2020.134917

Zhang, X. na, Guo, J. jun, Zou, X., Jin, D. chao, 2018. Pathogenic differences of the entomopathogenic fungus Isaria cateniannulata to the spider mite Tetranychus urticae (Trombidiformes: Tetranychidae) and its predator Euseius nicholsi (Mesostigmata: Phytoseiidae). Exp Appl Acarol. 10.1007/s10493-018-0247-x

